# Dedifferentiation alters chondrocyte nuclear mechanics during in vitro culture and expansion

**DOI:** 10.1101/2021.04.26.441500

**Authors:** Soham Ghosh, Adrienne K. Scott, Benjamin Seelbinder, Jeanne E. Barthold, Brittany M St. Martin, Samantha Kaonis, Stephanie E. Schneider, Jonathan T. Henderson, Corey P. Neu

## Abstract

Dedifferentiation of chondrocytes during *in vitro* passaging before implantation, and post implantation *in vivo*, is a critical limitation in cartilage tissue engineering. Several biophysical features define the dedifferentiated state including a flattened cell morphology and increased stress fiber formation. However, how dedifferentiation influences nuclear mechanics, and the possible long-term implications of this state, are unknown. In this study, we investigated how chondrocyte dedifferentiation affects the mechanics of the chromatin architecture inside the cell nucleus and the gene expression of the structural proteins located at the nuclear envelope. Through an experimental model of cell stretching and a detailed spatial intranuclear strain quantification, we identified that strain is amplified and distribution of strain within the chromatin is altered under tensile loading in the dedifferentiated state. Further, using a confocal microscopy image-based finite element model and simulation of cell stretching, we found that the cell shape is the primary determinant of the strain amplification inside the chondrocyte nucleus in the dedifferentiated state. Additionally, we found that nuclear envelope proteins have lower gene expression in the dedifferentiated state suggesting a weaker nuclear envelope which can further intensify the intranuclear strain amplification. Our results indicate that dedifferentiation and altered nuclear strain could promote gene expression changes at the nuclear envelope, thus promoting further deviation from chondrocyte phenotype. This study highlights the role of cell shape on nuclear mechanics and lays the groundwork to design biophysical strategies for the maintenance and enhancement of the chondrocyte phenotype during expansion with a goal of successful cartilage tissue engineering.

**SIGNIFICANCE:** Chondrocytes dedifferentiate into a fibroblast-like phenotype in a non-native biophysical environment. Using high resolution microscopy, intranuclear strain analysis, finite element method based computational modeling, and molecular biology techniques, we investigated how mechanical force causes abnormal intranuclear strain distribution in chondrocytes during the dedifferentiation process. Overall, our results suggest that the altered cell geometry aided by an altered or weakened nuclear envelope structure are responsible for abnormal intranuclear strain during chondrocyte dedifferentiation that can further deviate chondrocytes to a more dedifferentiated state.

## INTRODUCTION

Cartilage tissue engineering has received specific attention in the attempt to treat osteoarthritis, a debilitating disease affecting a large worldwide population (1). Pharmaceutical therapies to treat this degenerative disease are limited, and current disease management protocol relies on lifestyle changes and the use of drugs to ameliorate pain or to reduce inflammation. Tissue engineering and regenerative medicine are promising approaches with the goal to rebuild the damaged articular cartilage to restore the functions of the knee joint (2).

Autologous Chondrocyte Implantation (ACI) is a promising approach for cartilage tissue engineering, where chondrocytes extracted from the patient are expanded *in vitro* and then implanted back in the knee joint to regenerate functional articular cartilage (2–4). However, there are two critical challenges that limit the potential widespread application of this technology. First, from clinical follow up studies, it is now apparent that the success of the ACI treatment is limited because of post-operative fibrosis that leads to inferior biomechanical and functional response of the knee joint (5–9). The post-operative fibrosis is probably triggered by the dedifferentiation of the chondrocytes to fibroblast-like phenotype after implantation because the chondrocytes experience a mechanically fibrous and stiffer tissue microenvironment different than the native healthy cartilage environment, not conducive to the maintenance of the chondrocyte phenotype. The second critical challenge of ACI involves the expansion of the chondrocytes *in vitro*. During the expansion process and passaging, the chondrocytes dedifferentiate into fibroblast-like cells while cultured on a two-dimensional (2D) monolayer culture. Consequently, the scalability of the ACI technology is limited because growing enough cells for a large defect requires extensive time, which can further dedifferentiate chondrocytes in 2D culture.

Changes in mechanical loading of chondrocytes might lead to chondrocyte dedifferentiation *in vivo* and *in vitro*. Static and dynamic loading on the chondrocyte and its complex interaction with the pericellular matrix (PCM) and extracellular matrix (ECM) probably plays a critical mechanobiological role (10–13) in the dedifferentiation process (14). Mechanical stretching on chondrocytes in the native cartilage environment is shown to induce several pathophysiological responses mediated by several mechanobiological pathways (15–17). *In vitro*, the monolayer culture of chondrocytes on a dish imparts a passive two-dimensional mechanical stretch on the cell, aided by the flat cell shape and contractile forces imparted by F-actin stress fibers. Dedifferentiated chondrocytes in monolayer culture have a higher cell stiffness mediated by a cell membrane-F actin adhesion mechanism (18), indicating higher contractile forces within the cell. Interestingly, several studies indicate that if the cells are instead encapsulated in a three-dimensional (3D) environment, the chondrocyte phenotype prevails. Even the late passage chondroblasts (a term used to define a cell showing the phenotype of both chondrocyte and fibroblast) redifferentiate into chondrocytes (19, 20) when cultured in 3D, suggesting that releasing cell contraction experienced in 2D ameliorates the dedifferentiation.

The mechanical loading not only affects the chondrocyte cytoskeletal mechanics, but also it reaches and affects the cell nucleus as well (21–23) which might have unknown implications in the chondrocyte dedifferentiation process. The mechanics of the cell nucleus and its biological implication, termed as nuclear mechanobiology has recently been shown to be important in numerous biological contexts - in diseases (24, 25), gene expression (26), differentiation (27, 28), developmental biology (29) and tissue homeostasis (30). The study of nuclear mechanobiology investigates how the mechanical microenvironment (i.e., stretch, compression and shear) affects the spatiotemporal function of the nuclear envelope, the nucleoskeleton, the chromatin architecture, and hence the gene regulation. A recent study elucidated that the loss of extracellular matrix stiffness and the lack of three-dimensional chondrocyte matrix decreases the chondrocyte cell and nuclear stiffness, suggesting the critical role of cell mechanical environment in the chondrocyte phenotype maintenance and nuclear mechanics (31). Another recent study discovered that under shear stress, the chondrocyte nucleus experiences complex strain pattern resulted by heterogeneous intranuclear chromatin architecture, further highlighting the importance of chondrocyte nuclear mechanics (32). However, in the context of chondrocyte dedifferentiation it is not understood how the mechanical stretch and gradual dedifferentiation affects the intranuclear mechanics and the structural proteins of the nuclear envelope. Such knowledge could be potentially exploited for improving cartilage tissue engineering as intranuclear mechanics can directly affect the genomic expression stability of cells (33).

In this study, we hypothesized that cell spreading of chondrocytes during expansion processes results in altered cell mechanics leading to an abnormal strain burden in the nucleus via flat cell morphology and weaker nuclear envelope. To test this hypothesis, we mechanically stretched the chondrocytes in a controlled manner to visualize the strain anomaly in early passage and late passage chondrocyte nuclei using a technique already established before in the context of studying chondrocyte dedifferentiation and pathology (34–36). We found an amplified strain magnitude and an abnormal distribution of strain inside the chromatin of the chondrocyte that might be the cause of further degeneration of the chondrocyte phenotype, as indicated by changes in gene expression of the dedifferentiated chondrocytes compared to early passage chondrocytes. Next, using an image-based computational modeling of cell stretching we found that the flattened cell morphology is a key mediator of higher strain inside the cell nucleus. Gene expression studies of chondrocytes suggested a mechanically weaker nuclear envelope in later passages thus indicating an additional potential mechanism of amplified mechanical strain inside the cell nucleus at later passages.

## METHODS

### Chondrocyte extraction and culture

Primary bovine chondrocytes were harvested from the load-bearing region of the medial condyle from young bovine joints less than six months old (37, 38). The cells were seeded directly on the substrate after harvesting and these cells were described as Population Doubling 0 (PD0) cells, corresponding to passage 0. Harvested cells were cultured in DMEM-F12 with 10% FBS and passaged at around 80% confluency on petri dishes (Corning, 430165) until Population Doubling 8 (PD8), corresponding to passage 4.

### Imaging and geometrical characterization of chondrocytes

PD0 and PD8 chondrocytes were cultured on cell culture petri dishes (Corning), and StageFlexer Type I Collagen treated membranes (FlexCell International Corp) for 24 hours prior to fixation to allow for complete adhesion. Chondrocytes were fixed with cold 4% paraformaldehyde (Electron Microscopy Sciences) in 1X PBS, washed with 1X PBS three times, permeabilized in 0.1% Triton X-100 (Sigma Aldrich), and again washed with 1X PBS three times. Next, cells were incubated with DAPI (Thermo Fisher Scientific) for nuclear DNA staining (1:500) and Alexa Fluor 488 phalloidin (Thermo Fisher Scientific) for cytoplasmic F-actin staining (1:500). Fluorescent images of the PD0 and PD8 chondrocytes were captured using an inverted confocal microscope (Nikon Eclipse Ti AIR, Location, USA) with a 10× air objective and a 60× oil objective. Cells to image were chosen at random and distributed across the substrate. The area, the length and the aspect ratio of both the cell and the nucleus for imaged cell were measured using the image analysis software, Fiji.

### Gene expression analysis on PD0 and PD8 chondrocytes

Total RNA was extracted from the PD0 and PD8 chondrocytes from Type I Collagen treated membranes (FlexCell International Corp.) using AurumTM Total RNA Mini Kit, reverse transcribed into cDNA via iScriptTM Reverse Transcription Supermix. Real-time quantitative PCR was performed with SsoAdvancedTM Universal SYBR® Green Supermix in a CFX96 Touch™ thermocycler (all kits and devices from Bio-Rad Laboratories) using 10 ng of cDNA/ reaction. All data was normalized to the reference genes *HPRT1* and *RPL10A*. Primers were custom designed, and sequences are listed in **Supplementary Table 1**.

### Experimental set up for chondrocyte mechanical stretching

The FlexCell (FlexCell International Corp.) cell stretching device was used to apply controlled biaxial mechanical stretching on the chondrocytes. StageFlexer Type I Collagen treated membranes (FlexCell International Corp.) were used for the chondrocyte culture and stretching. The device was modified for custom imaging specific to the current application using a two-photon microscope (Olympus with a tunable Mai-Tai laser set at 540-600 nm). The first modification was to convert it from a vacuum device to a positive pressure device by placing a cap above the substrate **(Figure S1a)**. The second modification was to drill out the bottom support bracket under the loading post to allow for the objective to fit inside the device. The magnitude of the substrate stretch varied linearly with change in pressure, controlled by a pressure regulator programmed using a Labview interface **(Figure S1b-c)**. To calculate the substrate deformation, a previously developed technique was used (39). Briefly, the technique tracked the displacement of beads pre-embedded in the membrane during stretching. Subsequently, strain in the substrate was analyzed using the finite element method. Thus, the equibiaxial stretch mode on the substrate and on the cells was validated **(Figure S1d-e)**.

### Live chondrocyte stretching and imaging

Cells were incubated on the substrate for 24 hours to allow for complete adhesion prior to the stretching experiments. Nuclear DNA was stained with Hoechst 34580 (Life Technologies) before loading the substrate on the stretching device. The cells remained in the culture medium during imaging. Two photon microscopy (Olympus, 740 nm wavelength, 250 mW power for imaging the nucleus) with a 40× water objective was used to capture three images three minutes apart **(Figure S1b)** prior to stretching. The substrate, and in turn the chondrocytes, were then stretched. Next, three more images were captured after finding the chondrocytes in the new location. For each image capture the transmitted light was also used to visualize the overall cell shape.

### Spatial strain quantification inside the nucleus

Chondrocyte nucleus images captured before and after stretching were used to quantify spatial strain inside the nucleus with a technique called deformation microscopy (40, 41). Briefly, the image texture of the intranuclear space was utilized to achieve a high resolution intranuclear displacement map. Several strain measures including hydrostatic (related to volume change), deviatoric (related to shape change), and shear (angular deformation) were then computed from the displacement map. Hydrostatic strain was further categorized into tensile and compressive strain. A Hill’s function-based thresholding of the nuclear image was performed (42) to distinguish the heterochromatin (high chromatin density) and euchromatin (low chromatin density) domains.

### Image based modeling and simulation of strain in chondrocytes

A computational model was developed to quantify the influence of changes in cell geometry and mechanical properties on the deformation of nucleus at PD0 and PD8 chondrocytes. **Supplementary Note** explains the details of the image-based modeling and strain analysis. Briefly, PD0 and PD8 chondrocytes were cultured, fixed, and stained for DNA, F-actin and vinculin to image the nucleus, cytoplasm and focal adhesion complex respectively (**Figure 5a**). Three representative cells from each group were chosen at random for subsequent analysis. For each representative cell, the intracellular and intranuclear spaces were segmented to create three separate domains of cytoplasm, heterochromatin and euchromatin. Based on the confocal image stack, a three-dimensional geometry was created, meshed and imported into a finite element method software (COMSOL Multiphysics 5.3, Burlington, MA, USA). Further, appropriate governing equations and boundary condition were used to solve for the strain in the cytoplasm, heterochromatin and euchromatin.

### Effect of mechanical property on cell and nuclear strain: parametric study

A parametric study with the *in silico* cell model was used to investigate the effect of the cell mechanical properties on the intranuclear strain. For the parametric study, the elastic moduli of the cytoskeleton, euchromatin, and heterochromatin domains were varied for the PD0 and PD8 chondrocytes, while all other parameters (density and Poisson’s ratio) were held constant. Subsequently density and Poisson’s ratio were varied independently while other mechanical parameters were held constant. The values chosen for each parametric study are shown in **Supplementary Table 2**. The values assigned for each mechanical property are physiologically relevant and represent the extreme ends of the property value. The hydrostatic strain solution at each material node with the baseline mechanical properties (cytoplasm: Young’s modulus E = 500 Pa in PD0, and 900 Pa in PD8; euchromatin, E = 1000 Pa in PD0 and PD8; heterochromatin, E = 4000 Pa in PD0 and PD8), was compared against each new solution with a change in the parameter to assess how much the hydrostatic strain solutions differed when the parameters were varied.

### Statistical analysis

For cell and nuclear geometry analysis, *t*-test was performed with significance at p<0.01. For strain comparison between groups, *t*-test was performed with significance at p<0.05. For gene expression analysis study, *t*-test was performed with significance at p<0.01. For the analysis of the parametric studies pertaining to the computational model, slope *m* and coefficient of determination R^2^ was calculated using linear regression.

## RESULTS

### Cell and nuclear geometry and morphology

Early passage (PD0) chondrocytes displayed a round morphology with no visible F-actin stress fiber formation on either substrate: Type I Collagen treated membrane and the tissue culture plate (TCP) (**Figure 1a**). The nucleus was mostly located at the cell center. Late passage (PD8) cells, in contrast, showed a larger cell area, F-actin stress fiber formation, and the nucleus was often located off-center. Further quantification of the geometrical measures revealed that for all substrates the cell area increased by almost five times and the nuclear area increased by almost two times in PD8 cells compared to PD0 cells (**Figure 1b**). Additionally, the cellular and nuclear morphology of chondrocytes cultured on the Type I Collagen coated StageFlexer membranes showed similar phenotypes as they showed on tissue culture plate (TCP), which validated our approach of chondrocyte culture for the mechanical stretching experiment. Along with the cell geometry changes, the nuclear area compared to cell area, and the nuclear linear dimension compared to cell linear dimension decreased significantly in PD8 compared to PD0, a hallmark of deviation from the chondrocyte like phenotype to hypertrophic chondrocyte phenotype, as observed in native cartilages. In PD0, cell aspect ratio was maintained close to 1, signifying a round cell shape. In PD8, cell aspect ratio deviated significantly from 1, suggesting irregularly shaped cells. The nuclear aspect ratio showed a small deviation from the value 1 in PD8 compared to PD0 chondrocytes on collagen coated substrate, although the effect was not statistically significant. Irrespective of population doubling number, both the cell and nuclear area increased with the increasing stiffness of the substrate (stiffness of TCP > stiffness of Type I Collagen treated membrane), which was expected because a higher stiffness promotes larger cell and nuclear area through higher spreading of cells (43).

**Figure 1.**
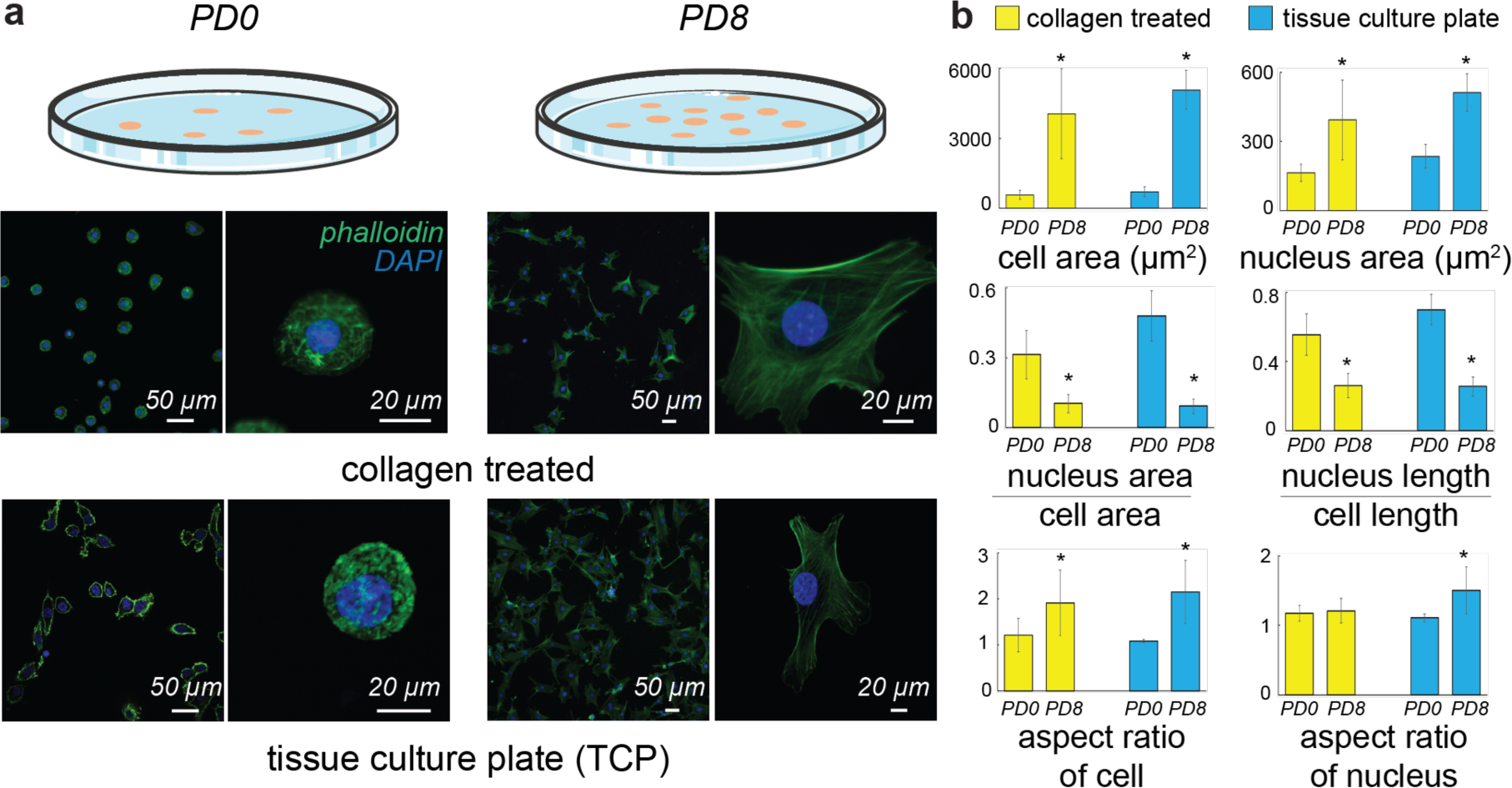
Dedifferentiation of chondrocytes in a monolayer culture was associated with a change in cell and nucleus phenotype. **(a)** Population Doubling 0 (PD0) – corresponding to passage 0 and Population Doubling 8 (PD8) – corresponding to passage 4 chondrocytes were stained for actin (phalloidin-GFP) and DNA (DAPI) to visualize cells and nuclei respectively using confocal microscopy at two different magnifications. Culture conditions included Type I collagen treated membranes and tissue culture plate (TCP). Irrespective of culture condition, the PD0 chondrocytes were smaller, rounder and displayed fewer actin stress fibers than the PD8 chondrocytes. **(b)** Several biophysical features were used to quantify the dedifferentiation mediated cell morphology changes between the PD0 and PD8. From PD0 to PD8, the cell and nuclear area increased, the nuclear dimensions increased compared to the cell dimensions, cells lost their round shape and approached more irregular shape; while nuclei maintain their round shape relatively higher compared to cells. *p < 0.01, number of cells analyzed > 60 from 3 different biological replicates.

### Passage specific gene expression of extracellular matrix proteins

Dedifferentiation of chondrocytes was associated with changes in extracellular matrix specific proteins. The gene expression of chondrocyte specific proteins (SOX9, ACAN, PRG4, COMP, COL2A1) decreased with dedifferentiation, while the fibroblast marker specific gene expression (COLA1, VIM, S100A4, Thy1) increased (**Figure 2**). Our gene expression data with young bovine articular cartilage chondrocyte agrees with previous reports of chondrocytes extracted from adult human knee articular cartilage (44), adult pig knee articular cartilage (45) and young bovine meniscus (46).

**Figure 2.**
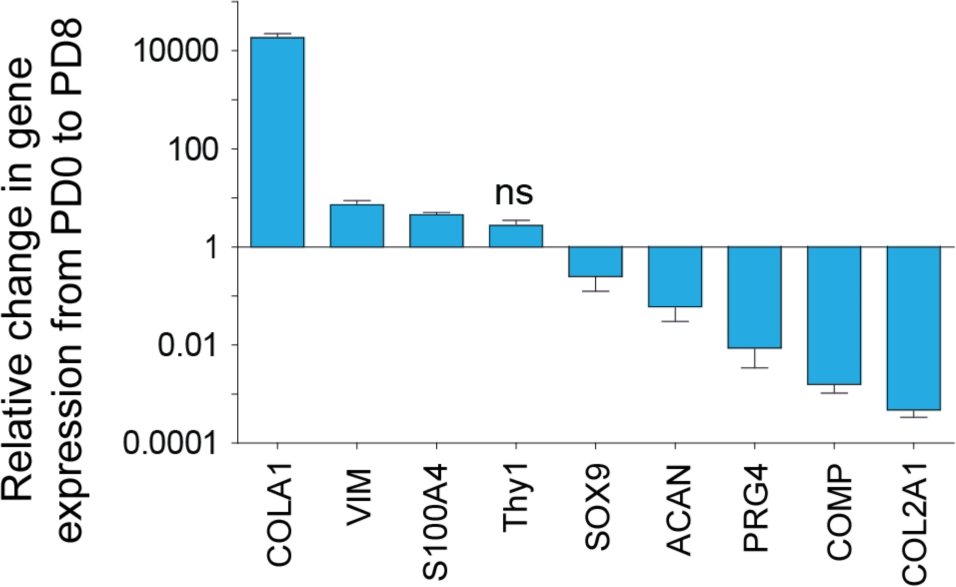
Dedifferentiation of chondrocytes in monolayer culture caused changes in extracellular matrix (ECM) related gene expression. Chondrocyte dedifferentiation from PD0 to PD8 was associated with an increase in the fibroblast specific ECM markers, and decrease in chondrocyte specific ECM markers. Except Thy1, all other gene expression changes were significant with p < 0.01, ns = not significant, number of samples = 4.

### Complex spatial intranuclear strain pattern

With the significant changes in cell morphology and gene expression between the PD0 and PD8 cell, we hypothesized that cellular geometry might influence strain transfer to the nucleus, contributing to mechanosensitive signaling that drives the dedifferentiation process. High resolution intranuclear spatial strain maps revealed an inhomogeneous deformation inside PD0 cell nuclei, while PD8 cell nuclei displayed a larger strain concentration at specific nuclear locations (**Figure 3a**). Shear and deviatoric strain values were comparable in PD0 and PD8. The hydrostatic strain values in the PD0 nuclei were lower overall (up to 0.35) compared to PD8 nuclei (up to 0.75), signifying that the PD8 nuclei displayed a magnified deformation for the same applied deformation on the cell surface. In PD0 cells, heterochromatin and euchromatin did not show any preferential strain localization. Alternatively, in PD8 cells high strain was associated with heterochromatin, which are pockets of high-density chromatin. The high strain regions in PD8 nuclei were most likely associated with local stress fibers in the cytoplasm, which was also predicted from the model as explained later (**Figure 5**). Most nuclear areas displayed a tensile strain, while compressive strain prevailed in some nuclear areas (**Figure 3b**). The histograms further showed, in detail, that the high chromatin density areas, i.e. the heterochromatin takes a more tensile strain burden compared to euchromatin in PD8 cells.

**Figure 3.**
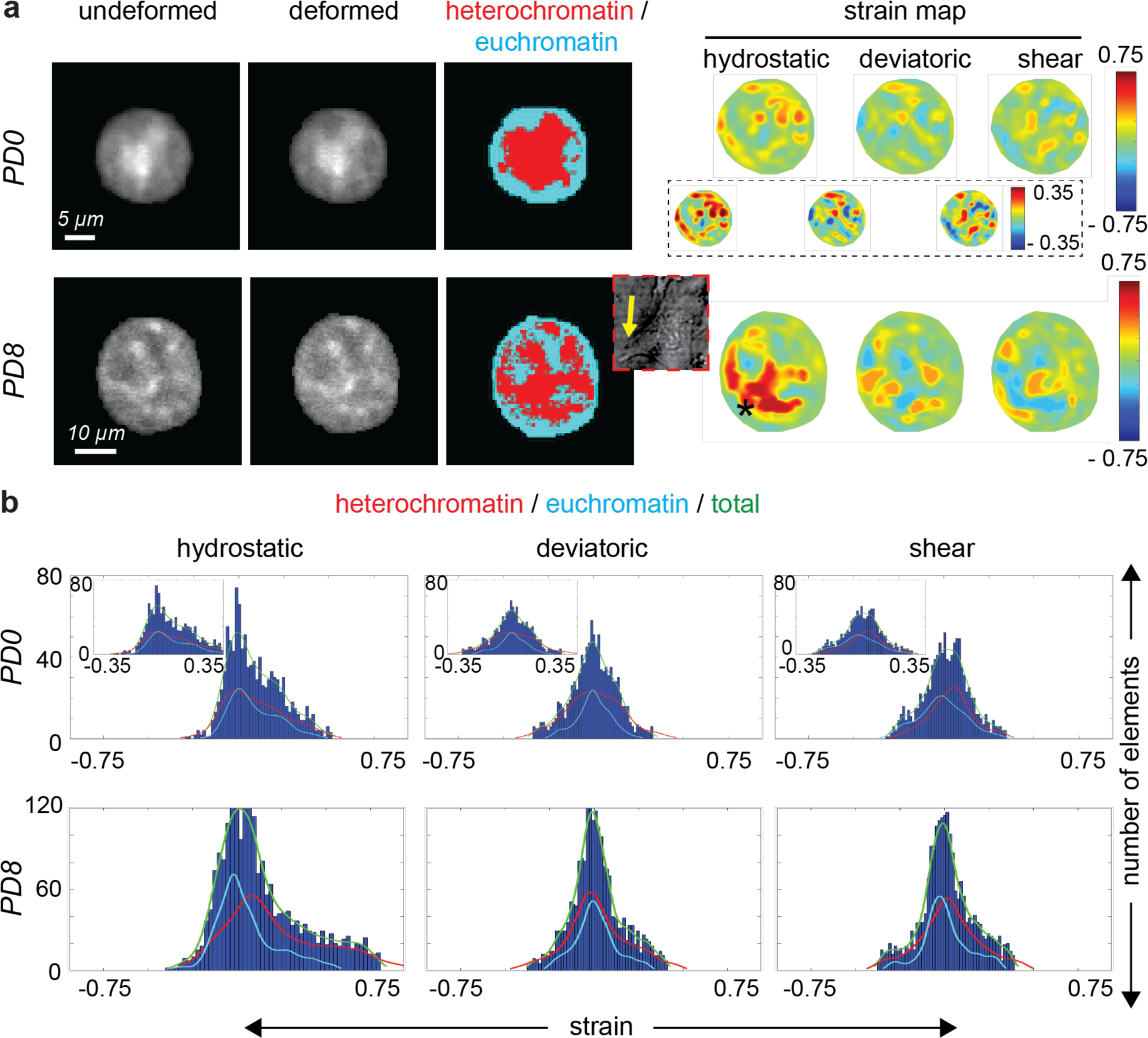
Passage dependent complex spatial strain patterns emerged inside the chondrocyte nucleus upon mechanical stretching of chondrocytes. **(a)** Using a FlexCell cell stretching device, a controlled equibiaxial stretch was applied to a collagen treated membrane culturing chondrocytes **(Figure S1).** Nuclei were imaged before stretching (undeformed) and at the stretched condition (deformed), followed by intranuclear spatial strain quantification using deformation microscopy, a technique developed previously (40). In PD0 cells, which were round shaped, magnitude of all the strain measures - hydrostatic (volume change related), deviatoric (shape change related) and shear were comparable with respect to each other. The high strain pockets were uniformly distributed in the nucleus irrespective of the chromatin density. Inset shows rescaled strain map in PD0 nucleus. In PD8 cells, which were irregular shaped, hydrostatic strain was significantly increased with respect to PD0 cells suggesting strain amplification, while deviatoric and shear strains are comparable in both PD0 and PD8 cell nuclei. High strain pockets (marked with star) were associated with the heterochromatin (high chromatin density regions), in the vicinity of cell protrusions known to be enriched with F-actin stress fibers. Cytoplasm image in the bright field is shown in the inset (not in the same scale as the nucleus images), with yellow arrow to show the cell protrusion adjacent to the high hydrostatic strain pocket. **(b)** Hydrostatic strain in the nucleus was mostly tensile, while deviatoric and shear strains followed mostly a normal distribution commensurate with the tensile cell stretching modality. Plots show histograms of the intranuclear elements for different strain measures for the representative nuclei in (a). In the PD0 nucleus, the heterochromatin (high density chromatin) and euchromatin (low density chromatin) strain distribution mostly overlapped, while in the PD8 nucleus the heterochromatin showed a higher tensile strain in most elements. Inset shows rescaled strain distribution for PD0 nucleus.

### Abnormal nuclear strain associated under tension in dedifferentiated cells

In dedifferentiated chondrocytes, the nucleus experienced an abnormal strain burden. In PD0 chondrocytes, the heterochromatin showed higher strain both in the tensile and the compressive modes (**Figure 4a**), suggesting that the heterochromatin might play a protective role to maintain a low strain burden in euchromatin. In PD8 chondrocytes, where the stiffer cell most likely imparted higher tension on the nuclear periphery, the heterochromatin experienced a significantly higher strain in tension and a lower strain in compression. Quantitative analysis (**Figure 4b**) further illustrated the strain differences as revealed by deformation microscopy. Thus, the protective effect of heterochromatin seemed to diminish in PD8 dedifferentiated cells.

**Figure 4.**
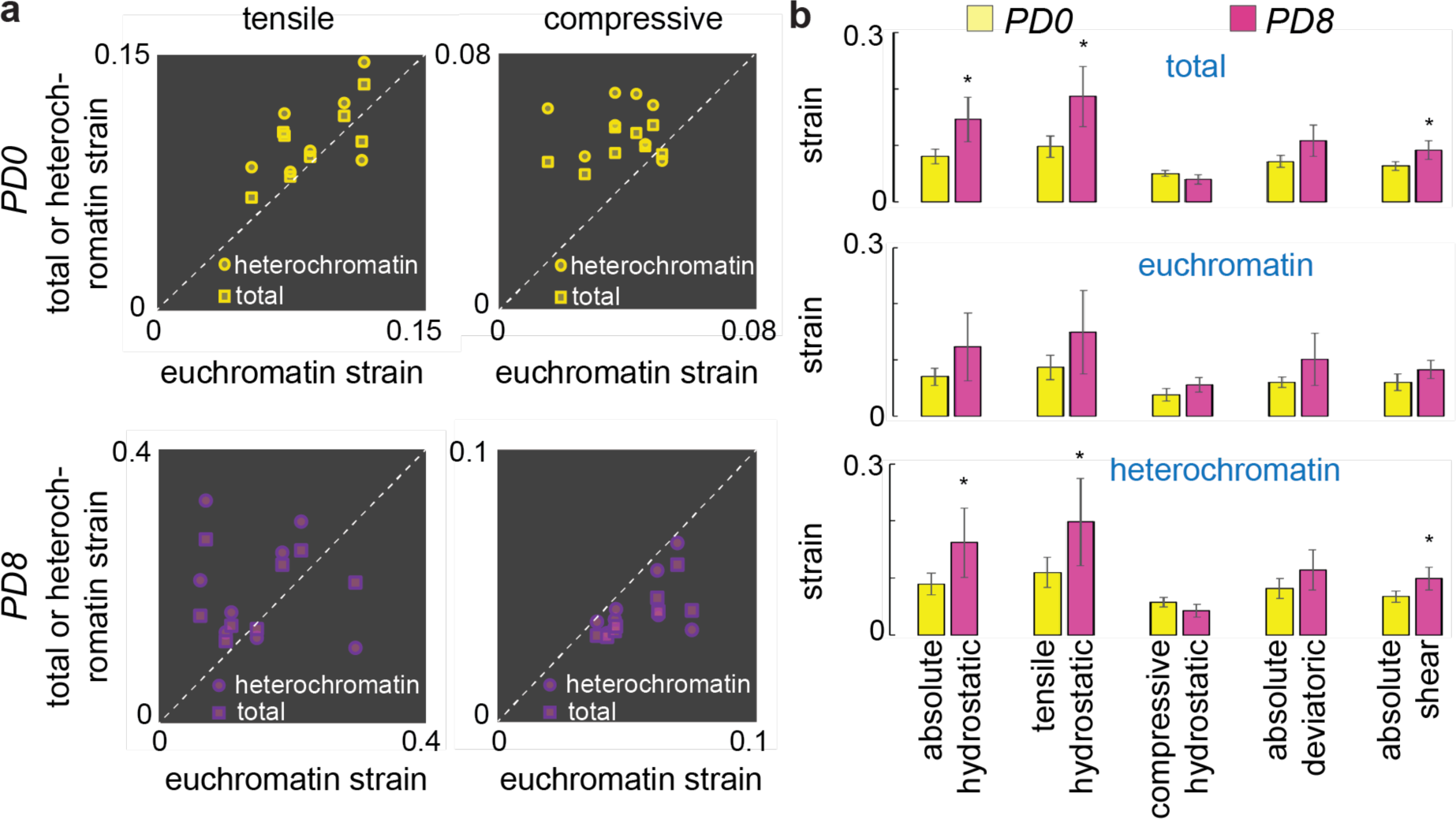
Dedifferentiation of chondrocytes was associated with abnormal intranuclear strain, most prominently visible in hydrostatic strain measure. **(a)** Each data point represents the average spatial strain for one specific nucleus. At PD0 chondrocytes, heterochromatin strain and total strain in the nucleus were slightly higher than the euchromatin strain both in the tensile and compressive modes. At PD8 chondrocytes, total strain and heterochromatin strain were lower compared to euchromatin strain in compressive mode, thus signifying a dense incompressible heterochromatin. In tensile mode, the heterochromatin experienced a higher strain compared to euchromatin. **(b)** Averaging the strain over multiple nuclei showed that the hydrostatic tensile strain was significantly higher in heterochromatin and in the nucleus as a whole compared to euchromatin, thus signifying an abnormal strain burden in the PD8 nuclei when compared to the PD0 nuclei. *p < 0.05, number of nuclei analyzed is at least 6 from 3 different biological replicates.

### Computational analysis of cell stretching

To understand why the PD8 chondrocyte nuclei experienced magnified strain, we performed the computational analysis of cell stretching and visualized the intranuclear strain due to the applied cell stretch. The computational analysis and the parametric study where the chondrocyte stretch was simulated, reveals that the intracellular and intranuclear strains are primarily influenced by the cell shape and geometry, while the mechanical properties of the cytoskeleton have a smaller effect on the nuclear strain. A lower 3D aspect ratio (maximum linear dimension parallel to the culture surface/ maximum cell height) of the cells promoted a lower strain in the nucleus as evident in the PD0 chondrocytes which were round in shape (**Figure 5a, Figure S1**). On the contrary, a larger 3D aspect ratio promoted a higher strain in the nucleus of PD8 chondrocytes which were flat in shape (**Figure 5a**, **Figure S1**). Volumetric hydrostatic strain maps showed key representative deformation features inside the PD0 and PD8 chondrocyte models (**Figure 5a, Figure S1**). The spatial hydrostatic strain distribution showed that on average, larger hydrostatic tensile strains were observed in the PD8 cell nucleus in comparison to the PD0 cell nucleus (**Figure 5b**). The higher strains in the nucleus were primarily associated with the cell protrusions, that were in the primary direction of tension propagation inside the cell (**Figure 5a**). These computational observations matched with the experimental observations, where some areas of the nucleus were associated with larger strains, likely due to the vicinity to the cell protrusions which were rich in F-actin stress fibers (**Figure 3a**). Overall larger hydrostatic tensile strains were observed in the PD8 cell nucleus in comparison to the PD0 cell nucleus from the computational studies (**Figure 5a, Figure 5b**), which agreed with the experimental measurement (**Figure 3**).

**Figure 5.**
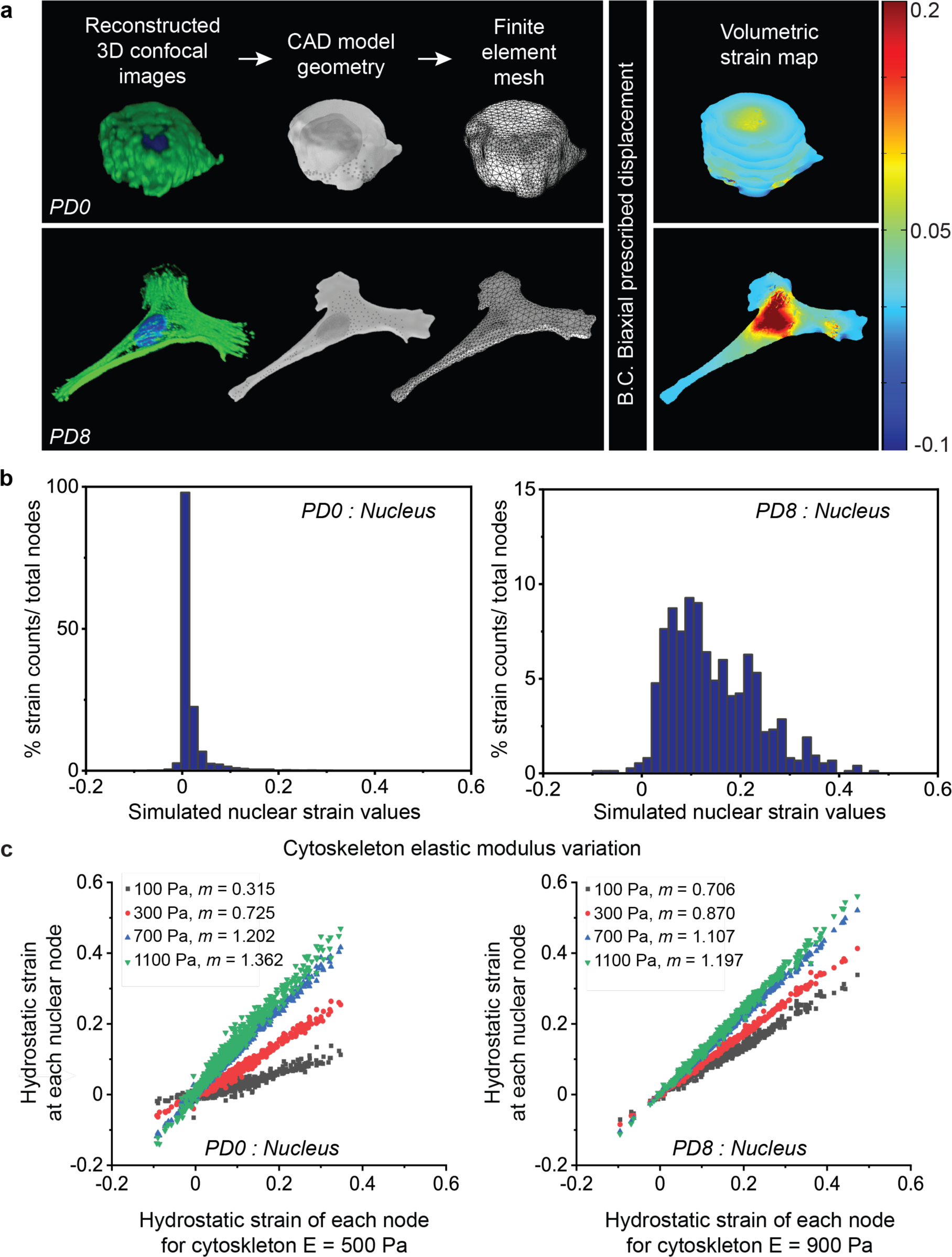
The chondrocyte shape change from PD0 to PD8 were a major factor determining the deformation of the cell and nucleus. **(a)** Confocal image stack of the cell and nucleus, along with the focal adhesion complex was used to create a three-dimensional model. PD8 cells were flattened with higher area and higher 3D aspect ratio, compared to PD0 cells which were round shaped with lower area and lower 3D aspect ratio. Appropriate governing equations and boundary conditions were used to analyze intranuclear and intracellular strain using finite element method. As the cell geometry altered from PD0 to PD8 (i.e., increased 3D aspect ratio and cell area), the tensile strain experienced by the nucleus almost doubled under the same boundary conditions, demonstrating the significant effect of the geometry change. **(b)** Histogram of the hydrostatic strain burden at the nuclear domain nodes. PD0 chondrocyte nuclei experienced lower value and lower range of strain, whereas PD8 chondrocyte nuclei experienced higher value and higher range of strain. **(c)** Cell morphology, not mechanical property was the key determinant of intranuclear strain. Irrespective of the population doubling status and geometry, the distribution of strain at computational nodes did not alter significantly with the variation of cytoskeleton elastic modulus (also see **Figure S1-S4**).

The parametric study showed that changing the mechanical properties had a less significant effect on the simulated strain of the system, when compared with the effect of cell shape and geometry. For example, as the elastic modulus of the cytoskeleton varied in comparison to the baseline initial values (E = 500 Pa for PD0 and E = 900 Pa for PD8), the strain pattern at each individual node in the nucleus remained similar as characterized by the R^2^ ∼ 0.98 in most cases **(Supplementary Table 3)**. The range of strain values changed with varying elastic modulus as characterized by deviation of slope *m* from 1, but overall, for the same elastic modulus (E = 100 Pa, 300 Pa, 700 Pa and 1100 Pa), the range of strain and maximum strain were higher in PD8 chondrocyte nuclei compared to PD0 chondrocyte nuclei (**Figure 5c**). The change in the heterochromatin and euchromatin elastic modulus had minor effect on the intranuclear strain distribution and values **(Figure S1, Supplementary Table 3)**. The varied Poisson’s ratio of cytoskeleton, euchromatin and heterochromatin also did not have a significant effect on the hydrostatic strain solutions at each nuclear node (**Figure S1, Supplementary Table 4**). However, the strain values in the nucleus for the PD0 and PD8 geometries varied by about a factor of two, as shown by the volumetric hydrostatic strain maps and histograms showing the distribution of strain (**Fig 5a, Figure 5b**). Therefore, changing the shape of the cell had the primary impact on the resulting nuclear strain range and magnitude compared to the material properties of the cell.

### Passage specific gene expression of nuclear envelope proteins

In addition to a change in geometry, we hypothesized that structural changes in the nuclear envelope could also lead to abnormal strain transfer to the chondrocyte nucleus. We found that dedifferentiation of chondrocytes was associated with a lower expression of several structural proteins in the nucleus (**Figure 6**), in addition to gene expression changes in extracellular matrix specific proteins (**Figure 2**). Expression of some key nuclear envelope related proteins such as Lamin B1, Lamin B2, Emerin and SUN 2 remained constant in the dedifferentiated state. Expression of some other proteins which provide stability to the nuclear envelope such as Lamin A/C, Nesprin 1, Nesprin 2, SUN 1 and Lemd3 decreased significantly. The highest decrease was associated with Lemd3 protein, which has a direct role in controlling the TGF-β and the BMP pathways, both critical factors in chondrogenic expression. SUN 1 is one of the two SUN proteins of the LINC (Linker of Nucleoskeleton and Cytoskeleton) complex which protrudes through the nuclear membrane, and its decreased expression severs the connection between the cytoskeleton and nucleoskeleton (25). Nesprin 1 and Nesprin 2 specifically bind to actin in cytoskeleton, and Lamin A/C creates a mesh-like structure under the inner nuclear membrane providing mechanical integrity to the cell nuclear envelope (25). Overall, this data suggested a weaker nuclear envelope, which in turn may amplify the intranuclear strain.

**Figure 6.**
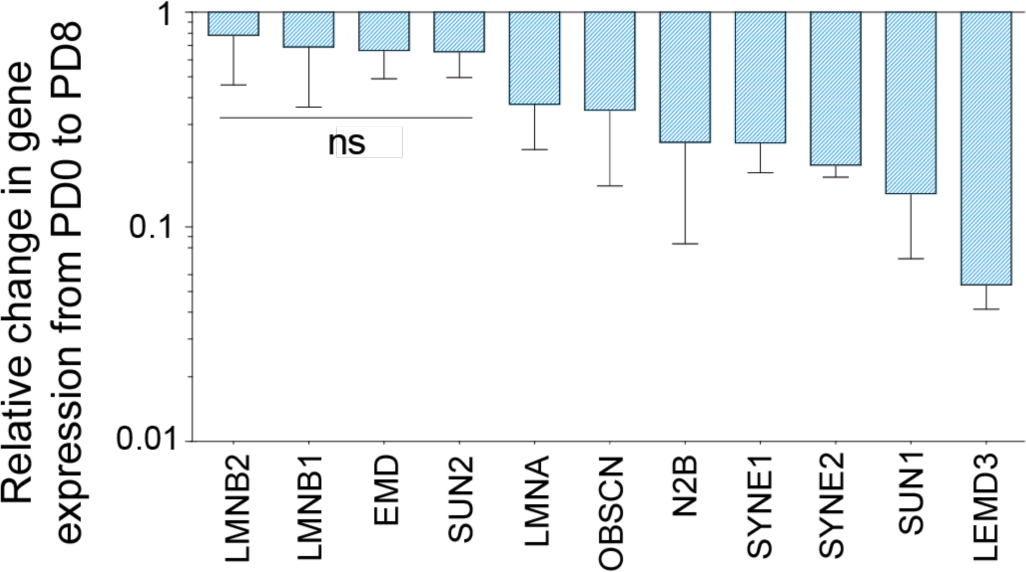
Dedifferentiation of chondrocyte in monolayer culture caused decrease in nuclear envelope related gene expression. Most nuclear envelope specific marker (e.g. Lamin A/C, nesprin 1 and 2 and SUN1) gene expression decreased in the dedifferentiated state. Significant (p < 0.01) gene expression change was observed in all cases except for the genes labeled with ns, ns = not significant, number of samples = 4.

## DISCUSSION

In this study, we investigated the mechanical behavior of early passage chondrocytes and dedifferentiated late passage chondrocytes, to elucidate the passage dependent gradual deviation from the chondrocyte phenotype. We found that the nuclear mechanics played a role behind this elusive chondrocyte behavior. The proposed mechanobiological model (**Figure 7**) summarizes possible contributing factors to the dedifferentiation process. Essentially, we hypothesize that the fate of chondrocytes is in a positive feedback loop where a chondrocyte is driven toward a fibroblast phenotype in two typical scenarios – after implantation *in vivo*, and before implantation during monolayer culture *in vitro*. In the proposed model, the mechanical stretch of the nucleus is a trigger, which can be induced by repeated biomechanical stretching *in vivo*, or during cell spreading and 2D cell shape maintenance *in vitro*. Utilizing a flattened cell shape and weaker nuclear envelope in dedifferentiated chondrocytes, the stretch causes amplified intranuclear strain and abnormal chromatin strain pattern thar are opposite to that of early passage chondrocyte. Such abnormal strain could shift the gene expression pattern to a more fibroblast specific lineage, which further increases the stretch on the nucleus, thus completing the positive feedback loop.

**Figure 7.**
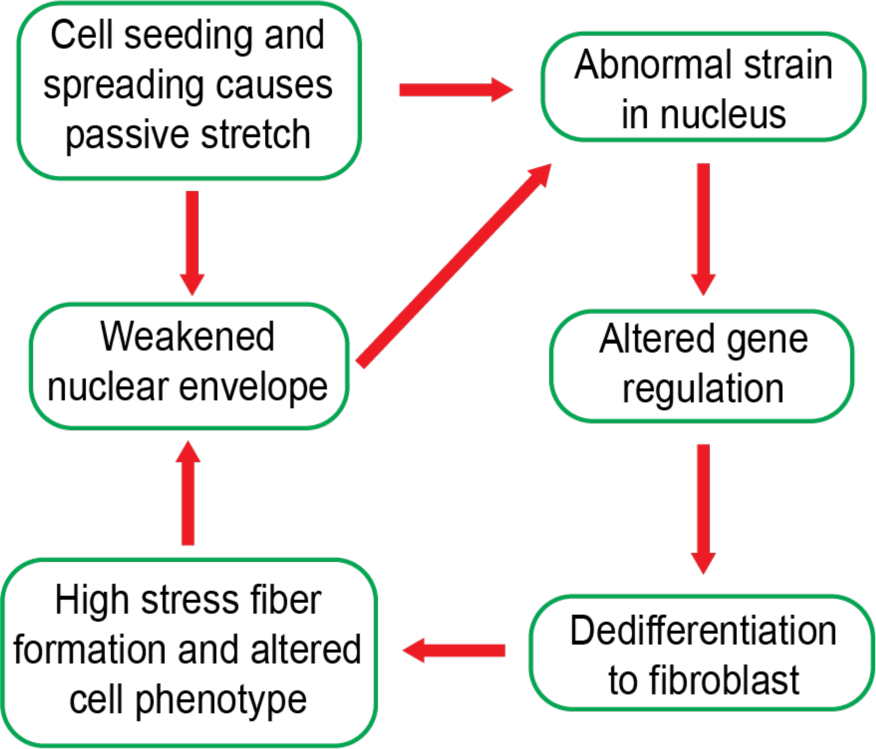
Proposed mechanobiological model summarizing the chondrocyte dedifferentiation in monolayer culture. A positive feedback loop driven by the stretch on the nucleus, abnormal intranuclear strain and gene dedifferentiation specific gene expression potentially favors the dedifferentiation dynamics.

Several studies thoroughly investigated what molecular changes happen with passage-driven dedifferentiation, and it was found that chondrogenic gene expression decreases and fibroblast specific expression increases with passaging (47, 48). A time lapse observation study was performed using chondrocyte specific reporter *Col11α2* to investigate whether the dedifferentiation was triggered by the chondrocyte division and proliferation, or by outgrowth of fibroblasts prevailing from the start of culture, but neither of these two mechanisms seemed to be the reason behind the dedifferentiation (49). Attempts have been previously made to understand the dedifferentiation mechanism, and subsequently redifferentiate the dedifferentiated cells to chondrocyte phenotype. Three dimensional encapsulation of late passage cells showed redifferentiation potential in several hydrogel systems such as agarose (50), polyethelene glycol and chondroitin sulfate (51), alginate (52, 53), dense collagen (20), and high density pellet culture (54, 55). There are several proposed mechanisms of how 3D culture promotes redifferentiation, such as the involvement of cytokines (56, 57) and microRNA (58). Our findings from this study suggests a biophysical mechanism in line with the existing reports. 3D culture decreases the stress fiber formation inside the cell and also imparts a round shape to the cell and nucleus (59). Therefore, in 3D, tension on the nucleus drastically reduces, changing the loading mode on the nucleus to compressive, potentially halting the dedifferentiation positive feedback loop proposed. Thus, the present study explains the redifferentiation mechanism in 3D. However, the 3D hydrogel-based therapy is not widely accepted for MACI procedure due to a lack of understanding the passaging limit of chondrocytes in the 2D environment and the inability of chondrocytes to substantially proliferate in the 3D environment (60).

Redifferentiation in 2D culture could be a promising approach as cells can be expanded to a large enough quantity, and their retrieval is straightforward. Several different approaches have been attempted to achieve this goal. Molecular biological interventions include the AIMP1 downregulation using siRNA to restore the TGF-β pathway (61), inhibition of BMP-2 using melanoma inhibitory activity protein (62) and upregulation of Sonic Hedgehog (Shh) pathway (63). Another approach is to induce stressed condition to the cells such as hypothermia (64), hypoxia (65), and applying intermittent hydrostatic pressure (66), but the underlying mechanism of stress-induced redifferentiation has not been completely explored. Importantly, disrupting the contractile function of the cytoskeleton, which in turn would change the deformation of the nucleus, is also a promising technique to prevent dedifferentiation (67, 68). Finally, epigenetic modifications of the dedifferentiated cells is a new approach on the horizon, where the chromatin architecture is directly targeted for redifferentiation (69). The outcome of our study which suggests a flattened cell, potentially weaker nuclear envelope suggested by the gene expression data, and amplified intranuclear strain pushes the chondrocyte phenotype toward a fibroblast; could provide biophysical design strategies to maintain chondrocyte phenotype in 2D culture. Potential approaches include overexpressing the nuclear envelope proteins while passaging the cells which can probably maintain the cells in chondrocyte phenotype with a round cell morphology. Additionally, it is also known the extent 2D cultured chondrocyte are able to redifferentiate when introduced into 3D culture after the expansion process, suggesting a possible ‘mechanical memory’ of the cells. This study lays the framework to further investigate how the mechanics gradually affects the chondrocyte phenotype possibly through a plastic and permanent change through intranuclear chromatin architecture, beyond which the chondrocyte memory is completely lost and therefore redifferentiaion becomes impossible.

Visualization of intranuclear strain in chondrocytes in the context of dedifferentiation and a weakened nucleus brings new insight of how the heterochromatin and euchromatin distributes the strain in a mechanically challenging condition. This finding can be of potential interest to the broad nuclear mechanobiology research, where research efforts have been made specific to chondrocytes (70) and mesenchymal stem cells with chondrogenic potential (27, 28, 71, 72) over recent years to understand the etiology of cartilage degeneration and regeneration.

Both the modeling and experimental results suggested that the chondrocyte 3D aspect ratio significantly affected the resulting intranuclear strain. In the computational model, as the chondrocyte 3D aspect ratio increased from PD0 to PD8, the tensile strain experienced by the nucleus increased under the same boundary conditions. The trend was consistent across a sample size of three cells for both PD0 and PD8. This trend confirmed the experimental data that indicated an overall increase in hydrostatic tensile strain found in the nucleus of the PD8 chondrocytes in comparison to the PD0 chondrocytes. Although major trends were confirmed by the computational model, the exact values of strain reported by the model would not be the same as the experimental results. These inconsistencies were expected due to the simplicity of the isotropic linear elastic model in the computational framework. The parametric study demonstrated that a change in the mechanical properties in the computational model did not have a large effect on the solution of stress and strain at each material node. Therefore, the cell and nuclear geometry changed from PD0 and PD8 was a major factor determining the abnormal increase in tensile strain in the nucleus of the PD8 cell. For example, even if the cytoskeleton stiffness increased in both the PD0 to PD8 cell, the predominate factor that caused an increased nuclear strain was the change in cell geometry toward a flattened shape as the chondrocyte dedifferentiated.

## CONCLUSION

In conclusion, we found that the flattened cell shape and potentially weaker nuclear envelope of expanded chondrocytes leads to an induced abnormal intranuclear strain. We propose that this abnormal nuclear stain could be a major factor driving chondrocyte dedifferentiation to a fibrotic phenotype of cells in monolayer culture, and potentially in mechanically challenged native cartilage after implantation. Abnormal intranuclear strain and weaker nuclear envelope are the key components of the proposed mechanobiological circuit that works in a positive feedback loop to deviate the chondrocyte from its original phenotype. The findings of this study have the potential to advance the autologous chondrocyte implantation (ACI) procedure by rational, biophysics-driven tissue engineering design strategy.

## SUPPORTING MATERIAL

Supporting material can be found online at Biophysical Journal Website

## AUTHOR CONTRIBUTIONS

S.G., B.S., and C.P.N. conceptualized the study, J.H., B.S., J.E.B., B.S.M.M., A.K.S., S.E.S., and S.G. performed the experiments, S.G. analyzed the data with the contribution from S.K.; A.K.S. performed the simulation and the computational study, S.G wrote the manuscript with contribution from all other authors, C.P.N. provided the funding and resources.

## ACKNOWLEDGEMENTS

The authors would like to acknowledge funding from NIH R01 AR063712, NIH R21 AR064178, NIH R21 AR066230, NSF CAREER1349735.

## CONFLICT OF INTEREST

The authors declare no conflict of interest.

## SUPPORTING MATERIALS

**Supplementary Table 1.**
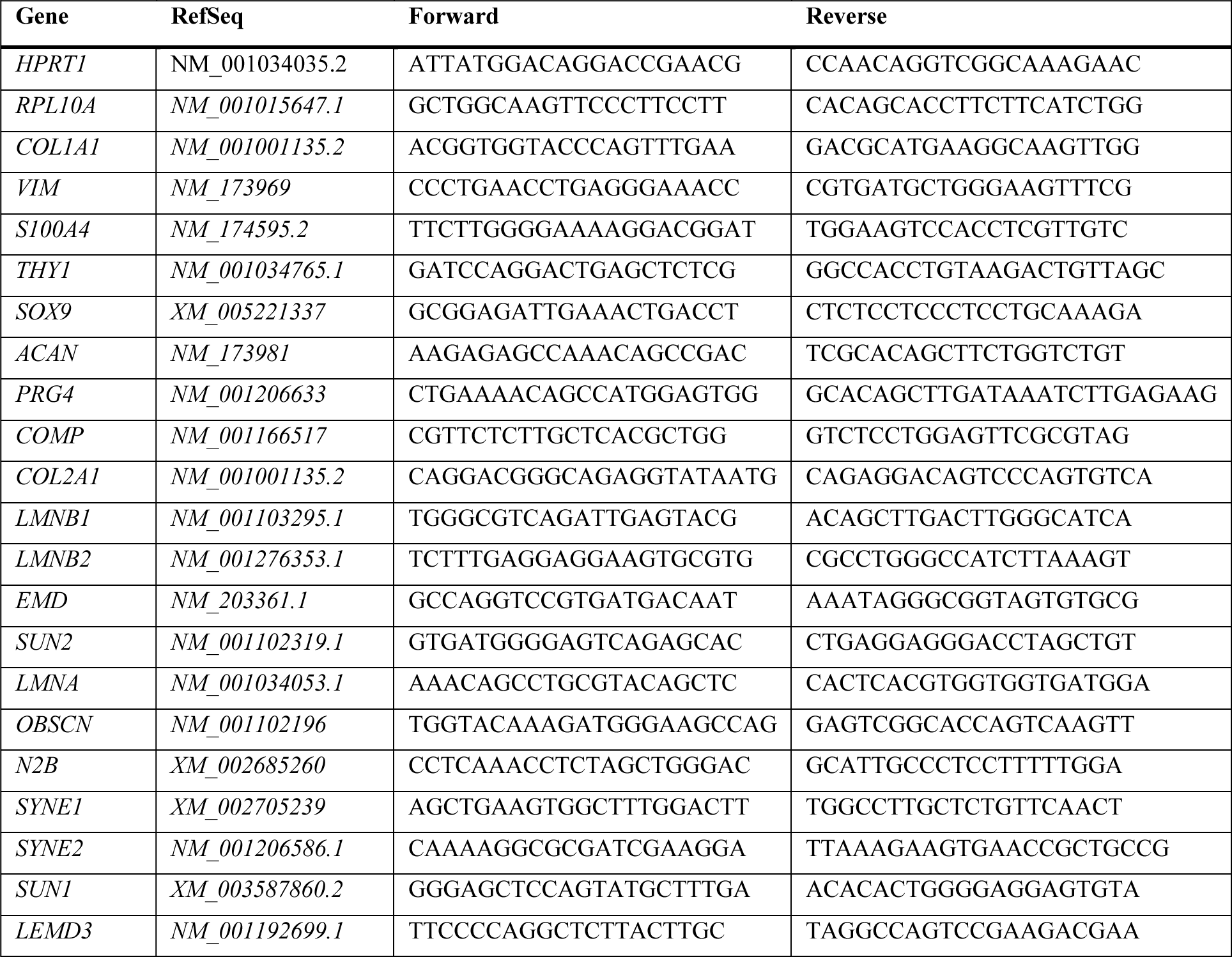
PCR Primer Sequences

**Supplementary Table 2.** Mechanical properties assigned to the computational domains for parametric study

|  | Cytoskeleton<br>Domain | Euchromatin<br>Domain | Heterochromatin<br>Domain |
| --- | --- | --- | --- |
| Elastic Modulus (Pa) | 100-1300 | 600-1400 | 2000-10000 |
| Poisson's Ratio | 0.2-0.45 | 0.2-0.45 | 0.2-0.45 |
| Density (kg/m<sup>3</sup>) | 1000-1400 | 1000-1400 | 1000-1400 |

**Supplementary Table 3.** Summary of results with parametric study - elastic modulus variation

**Cytoskeleton elastic modulus variation**
| *PD0: Baseline $E = 500$ Pa* |  |  | *PD8: Baseline $E = 900$ Pa* |  |  |
| --- | --- | --- | --- | --- | --- |
| | $m$ | $R^2$ | | $m$ | $R^2$ |
| $E = 100$ Pa | 0.315 | 0.938 | $E = 500$ Pa | 0.706 | 0.985 |
| $E = 300$ Pa | 0.725 | 0.990 | $E = 700$ Pa | 0.870 | 0.997 |
| $E = 700$ Pa | 1.202 | 0.995 | $E = 1100$ Pa | 1.107 | 0.999 |
| $E = 1100$ Pa | 1.362 | 0.984 | $E = 1300$ Pa | 1.197 | 0.995 |

**Euchromatin elastic modulus variation**
| *PD0: Baseline $E = 1000$ Pa* |  |  | *PD8: Baseline $E = 1000$ Pa* |  |  |
| --- | --- | --- | --- | --- | --- |
| | $m$ | $R^2$ | | $m$ | $R^2$ |
| $E = 600$ Pa | 1.327 | 0.959 | $E = 600$ Pa | 1.290 | 0.966 |
| $E = 800$ Pa | 1.133 | 0.992 | $E = 800$ Pa | 1.125 | 0.993 |
| $E = 1200$ Pa | 0.902 | 0.994 | $E = 1200$ Pa | 0.901 | 0.995 |
| $E = 1400$ Pa | 0.826 | 0.980 | $E = 1400$ Pa | 0.821 | 0.980 |

**Heterochromatin elastic modulus variation**
| *PD0: Baseline $E = 4000$ Pa* |  |  | *PD8: Baseline $E = 4000$ Pa* |  |  |
| --- | --- | --- | --- | --- | --- |
| | $m$ | $R^2$ | | $m$ | $R^2$ |
| $E = 2000$ Pa | 1.007 | 0.941 | $E = 2000$ Pa | 0.953 | 0.898 |
| $E = 3000$ Pa | 1.004 | 0.991 | $E = 3000$ Pa | 0.988 | 0.984 |
| $E = 5000$ Pa | 0.996 | 0.996 | $E = 5000$ Pa | 1.002 | 0.991 |
| $E = 10000$ Pa | 1.002 | 0.994 | $E = 10000$ Pa | 0.979 | 0.883 |

**Supplementary Table 4.** Summary of Summary of results with parametric study - Poisson’s ratio variation

**Cytoskeleton Poisson's ratio variation**
| *PD0: Baseline $\nu = 0.3$* |  |  | *PD8: Baseline $\nu = 0.3$* |  |  |
| --- | --- | --- | --- | --- | --- |
| | *m* | $R^2$ | | *m* | $R^2$ |
| $\nu = 0.2$ | 0.950 | 0.997 | $\nu = 0.2$ | 0.959 | 0.998 |
| $\nu = 0.25$ | 0.969 | 0.999 | $\nu = 0.25$ | 0.974 | 0.999 |
| $\nu = 0.35$ | 1.049 | 0.999 | $\nu = 0.35$ | 1.042 | 0.999 |
| $\nu = 0.4$ | 1.129 | 0.992 | $\nu = 0.4$ | 1.113 | 0.995 |
| $\nu = 0.45$ | 1.283 | 0.970 | $\nu = 0.45$ | 1.251 | 0.983 |

**Euchromatin Poisson's ratio variation**
| *PD0: Baseline $\nu = 0.3$* |  |  | *PD8: Baseline $\nu = 0.3$* |  |  |
| --- | --- | --- | --- | --- | --- |
| | *m* | $R^2$ | | *m* | $R^2$ |
| $\nu = 0.2$ | 1.249 | 0.994 | $\nu = 0.2$ | 1.206 | 0.988 |
| $\nu = 0.25$ | 1.136 | 0.998 | $\nu = 0.25$ | 1.116 | 0.996 |
| $\nu = 0.35$ | 0.834 | 0.994 | $\nu = 0.35$ | 0.848 | 0.989 |
| $\nu = 0.4$ | 0.631 | 0.954 | $\nu = 0.4$ | 0.641 | 0.912 |
| $\nu = 0.45$ | 0.379 | 0.728 | $\nu = 0.45$ | 0.346 | 0.508 |

**Heterochromatin Poisson's ratio variation**
| *PD0: Baseline $\nu = 0.3$* |  |  | *PD8: Baseline $\nu = 0.3$* |  |  |
| --- | --- | --- | --- | --- | --- |
| | *m* | $R^2$ | | *m* | $R^2$ |
| $\nu = 0.2$ | 1.028 | 0.983 | $\nu = 0.2$ | 0.944 | 0.957 |
| $\nu = 0.25$ | 1.015 | 0.995 | $\nu = 0.25$ | 0.971 | 0.989 |
| $\nu = 0.35$ | 0.984 | 0.994 | $\nu = 0.35$ | 1.030 | 0.988 |
| $\nu = 0.4$ | 0.965 | 0.974 | $\nu = 0.4$ | 1.062 | 0.951 |
| $\nu = 0.45$ | 0.945 | 0.936 | $\nu = 0.45$ | 1.095 | 0.891 |

## Supplementary Note: Simulation of Intranuclear Strain in Chondrocytes

### Immunostaining of chondrocytes and imaging

Population Doubling (PD0) corresponding to passage 0 and Population Doubling (PD8) corresponding to passage 4 chondrocytes were cultured, stained and imaged to generate the image-based model for further analysis. Chondrocytes were incubated on the Type I collagen coated substrate for 24 hours to allow for complete adhesion. Next, chondrocytes were fixed with cold 4% paraformaldehyde (Thermo Fisher Scientific) in 1X PBS, permeabilized in 0.1% Triton X-100 (Sigma Aldrich), and blocked with blocking buffer consisting of 1% bovine serum albumin (Sigma Aldrich), 10% Natural Goat Serum (Sigma Aldrich), 0.3% Tween 20 (Sigma Aldrich) in 1X PBS. Next, samples were incubated at 4°C overnight with an anti-vinculin primary monoclonal antibody (Sigma Aldrich) in antibody buffer (1% bovine serum albumin and 0.1% Tween in 1X PBS, at 1:50 dilution). Following primary incubation, chondrocytes were washed, and incubated with a secondary antibody, Alexa 546 anti-mouse, at 1:200 dilution (Thermo Fisher Scientific). To visualize the nucleus/DNA and cytoskeleton F-actin, chondrocytes were stained with Hoechst 34580 (Life Technologies) and Alexa Fluor 488 phalloidin (Thermo Fisher Scientific).

High magnification fluorescent *z*-stack images of the PD0 and PD8 chondrocyte chondrocytes were captured using an inverted confocal microscope (Nikon Eclipse Ti AIR, Location, USA) with a 60× oil objective. Thirty z-planes were taken for each cell with a step size that ranged from 0.2 µm to 0.3µm. Cells to image were chosen at random, distributed across the dish.

### Image based modeling of the chondrocyte cytoplasm and intranuclear space

From the z-stack of confocal microscope images, a three-dimensional model of both PD0 and PD8 chondrocytes were reconstructed. A custom MATLAB (The Mathworks Inc., Natick, MA, USA) program was used to process the z-stack images of the nucleus, cytoplasm consisting F-actin and cell surface focal adhesion complex containing vinculin. Using these three markers, each cell was segmented into three domains: cytoplasm, euchromatin and heterochromatin. The chromatin DNA in the nucleus, imaged in the blue channel, was thresholded to define the euchromatin and the heterochromatin regions, using the same technique described in the main text. The filamentous actin staining (green channel) was used to determine the cytoplasm domain. Based on the segmented regions, sterolithography (STL) files were created containing the triangulated surface information for each domain. These sterolithography files were then imported into open-source mesh processing system MeshLab (CNP, Rome, Italy) to further smooth the geometry (1). The smoothed geometries could then be imported as STL files into a finite element software (COMSOL Multiphysics 5.3, Burlington, MA, USA). Using a custom MATLAB code, the coordinates of the vinculin location points with respect to the cell were found using the confocal images and mapped onto the three-dimensional model using COMSOL with MATLAB interphase.

### Governing equations and boundary conditions

After importing the 3D geometrical model in COMSOL, the cell was divided into three computational domains: cytoplasm, euchromatin and heterochromatin. An equibiaxial prescribed displacement boundary condition of 5 µm was imposed on the bottom surface of the cell where the vinculin points were tracked and plotted as described above. All other boundary conditions were set as free, meaning that no loads or constraints acted on the boundaries and these boundaries were traction free.

Assuming linear elastic and isotropic materials for all three domains, the following governing equations were used to solve for the stationary solution of stress and strain with the applied boundary conditions previously described.

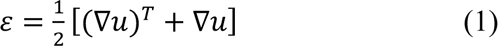

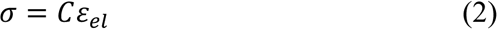

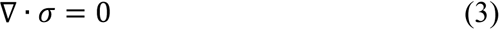

Equation (1) is the kinematic relation used to solve strain, *ε*, with the displacements, *u*. The total stress, *σ,* was calculated using Hooke’s law (Equation (2)), where *C* is the stiffness matrix and *ε_el_* is the elastic strain. Additionally, equilibrium equation with negligible body forces, Equation (3), was applied.

### Computational Procedure

All STL files were imported as a mesh into a finite element solver COMSOL Multiphysics 5.3 to create the three-dimensional solid geometry of the three domains. The material properties were assumed as follows (2): cytoplasm, Young’s modulus E = 500 Pa in PD0, and 900 Pa in PD8; heterochromatin, E = 4000 Pa in PD0 and PD8; euchromatin, E = 1000 Pa in PD0 and PD8. In all cases Poisson’s ratio was assumed to be 0.3 and density was kept constant at 1200 kg/m^3^. For each computational domain, a fine tetrahedral meshing scheme was used. Direct PARDISO solver was used to solve for stress and strain of the computational domains. With the known calculated strain values, the hydrostatic strain was found for each node with the following equation:

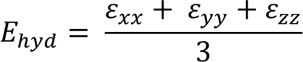

## Supplementary Figures

**Figure S1.**
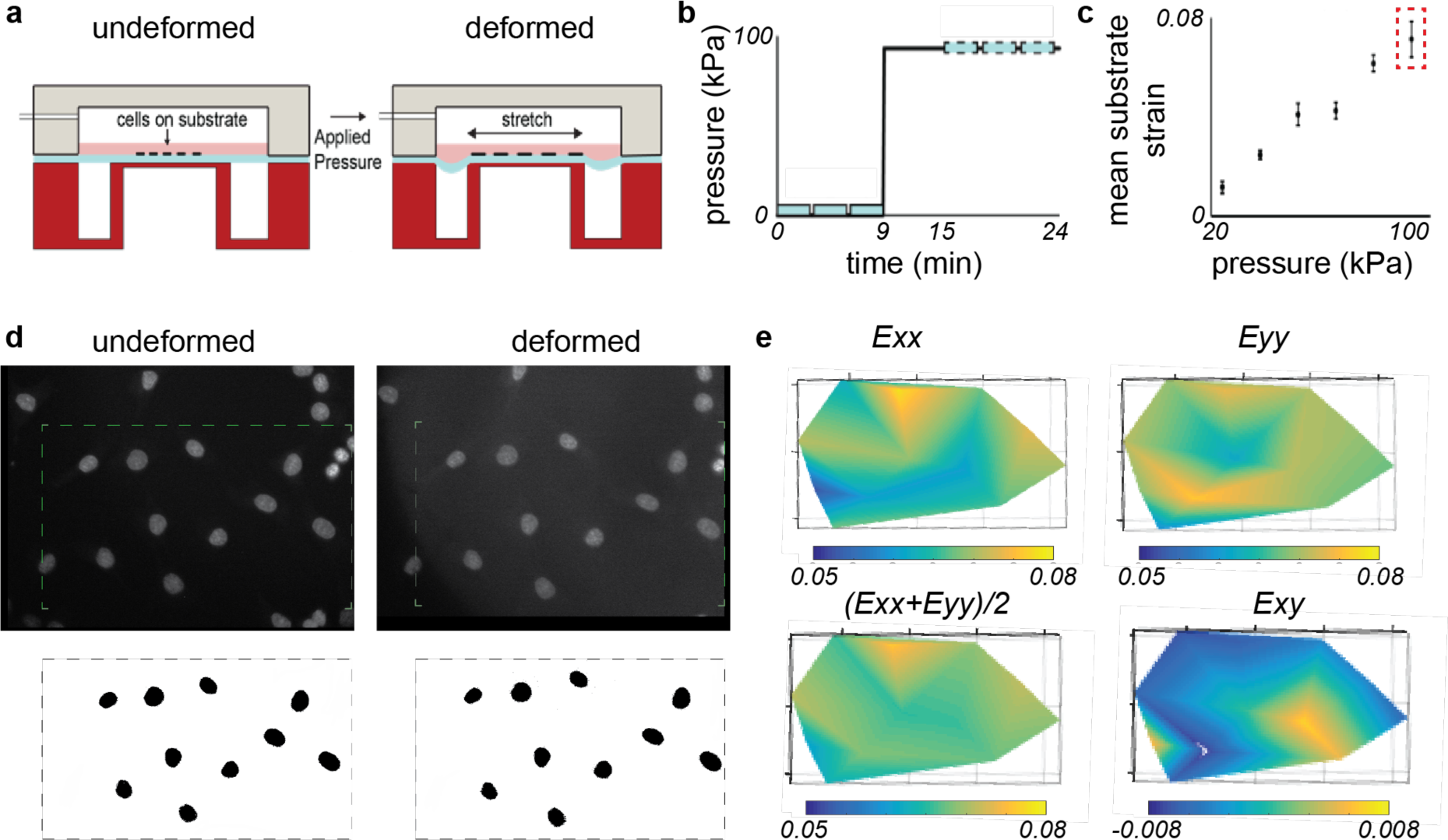
Equibiaxial mechanical stretching set up for the membrane with validation of deformation. **(a)** FlexCell stretching device allowed the chondrocytes to be seeded and stretched using a controlled external pneumatic pressure, while the cell nucleus could be imaged using a two-photon microscope. **(b)** Imaging of the nucleus was performed every 3 minutes at the undeformed and deformed states of the chondrocytes. The deformation of chondrocytes was induced by the stretching of membrane using external pneumatic pressure. **(c)** Mean substrate strain with increasing pressure showing the calibration of the cell stretching set up. Red dotted rectangle signifies the pressure (100 kPa) that was used for all experiments in this study. **(d)** The nuclei moved with respect to each other while the membrane stretched as shown by the undeformed and deformed membrane images (top panel). A thresholding was done to track the centroid position of each individual nuclei (bottom panel). **(e)** Characterization of the membrane stretch quantified by the strain (3). The *Exy* strain was an order of magnitude lower than the *Exx* and *Exy* strain, confirming a nearly equibiaxial stretch in the membrane.

**Figure S2.**
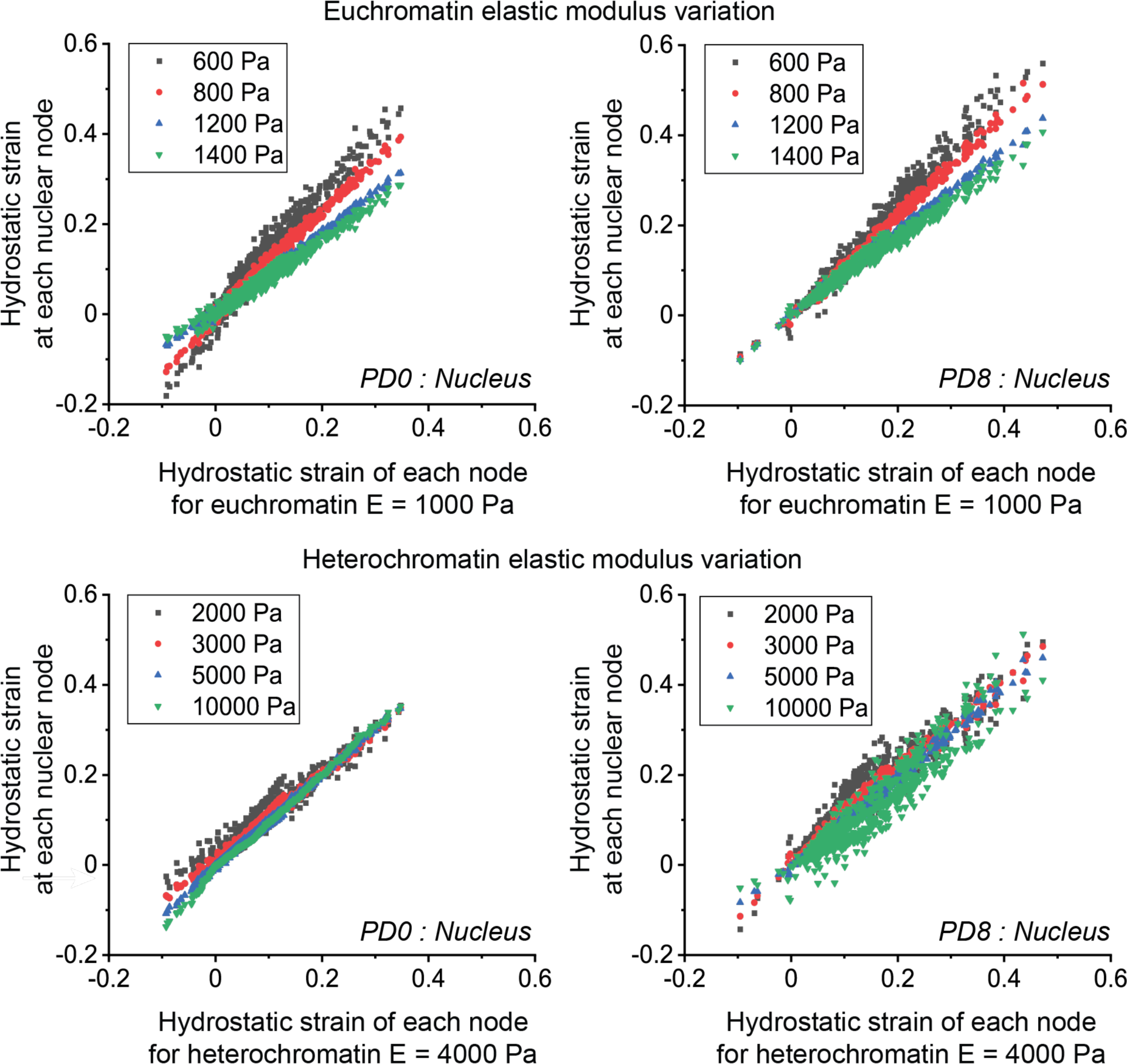
Effect of the euchromatin and heterochromatin elastic moduli on intranuclear deformation. Parametric study showed that geometry (early passage vs late passage) determined the range of strain in chondrocyte nucleus, even when the euchromatin and heterochromatin elastic moduli were varied.

**Figure S3.**
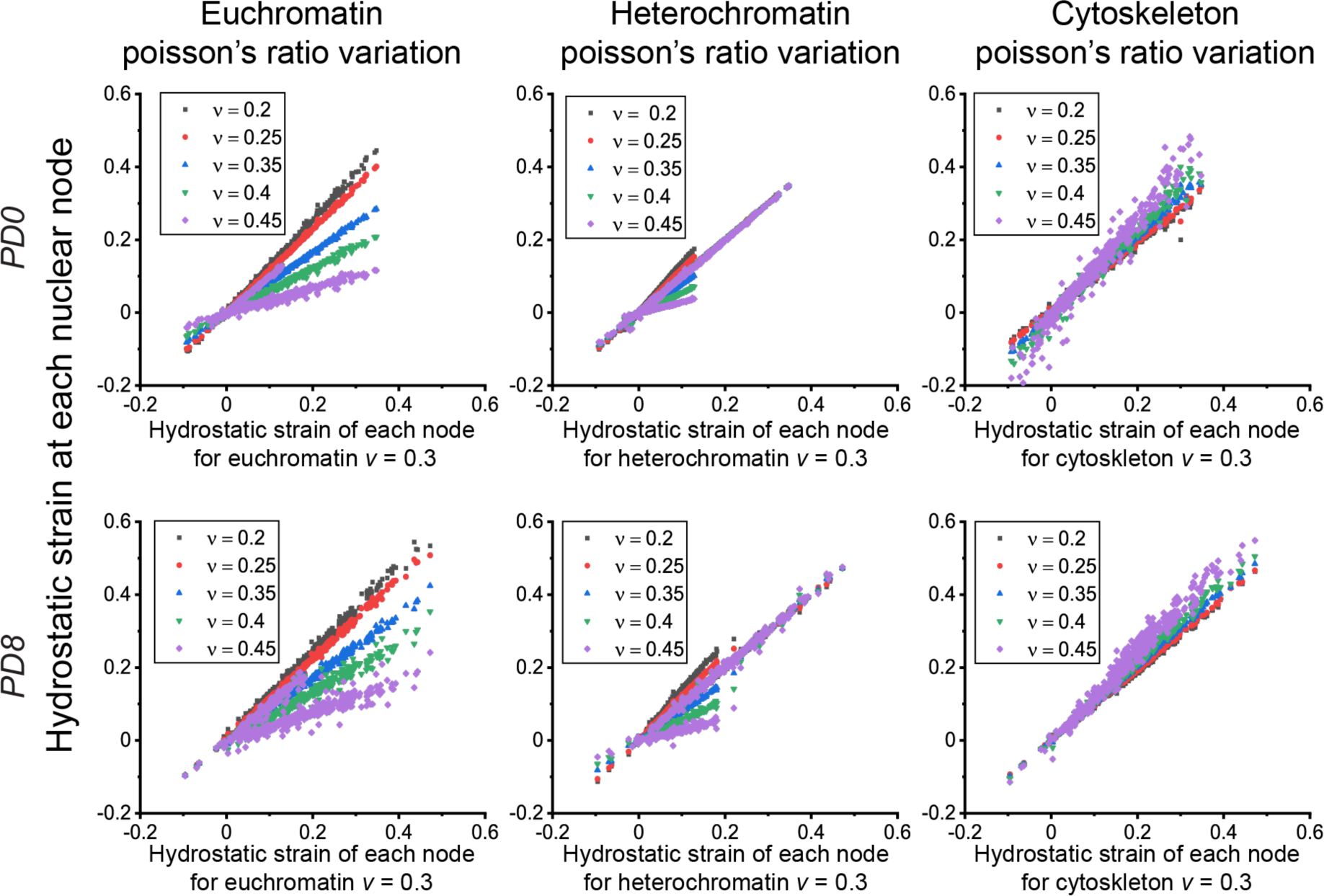
Effect of the cell domain Poisson’s ratio on deformation. Parametric study showed that geometry (early passage vs late passage) determined the range of strain in chondrocyte nucleus, even when the Poisson’s ratio was varied in cytoskeleton, euchromatin and heterochromatin.

**Figure S4.**
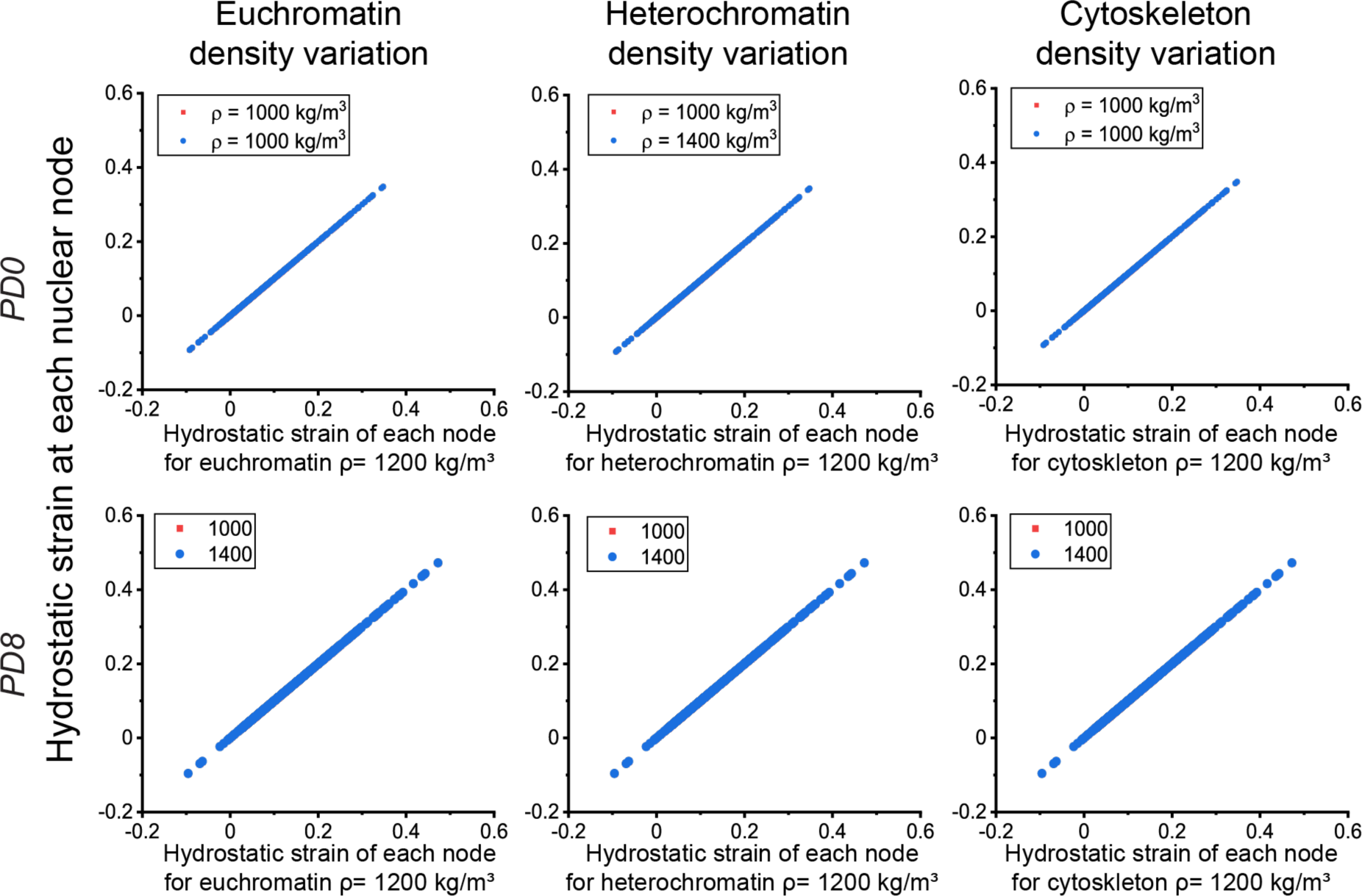
Effect of the cell domain densities on deformation. Parametric study showed that the density of the cytoskeleton, euchromatin and heterochromatin had no effect on the intranuclear deformation (*m* = 1 and R^2^ = 1 in all cases). The reason is the material model we used for the chondrocyte deformation simulation had no density component.

**Figure S5.**
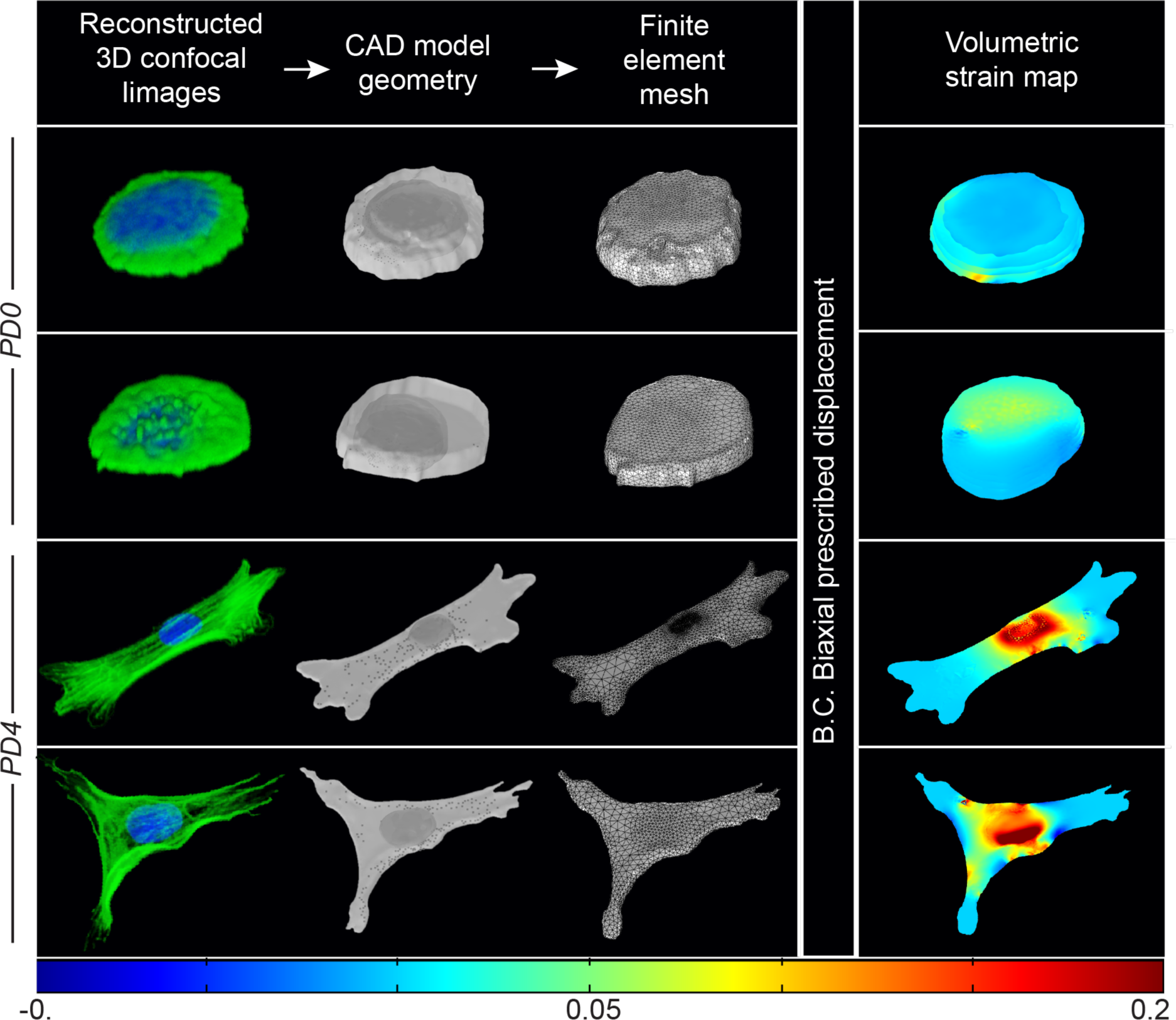
Additional representative examples of PD0 (early passage) and PD8 (late passage) chondrocytes displaying the simulated intracellular and intranuclear strain.

## Notes

### Competing Interest Statement

The authors have declared no competing interest.

